# Macrophages reshape cytokine responses and bacterial spatial organization in an airway epithelial cell culture model

**DOI:** 10.64898/2026.05.27.727787

**Authors:** Alexander F. Melanson, Jenny J. Persson, Søren Molin, Helle Krogh Johansen

## Abstract

The increasing prevalence of antibiotic-resistant bacterial infections highlights the need for physiologically relevant *in vitro* models that recapitulate host–pathogen interactions. *Pseudomonas aeruginosa* is a clinically important opportunistic pathogen associated with hospital-acquired infections and chronic airway diseases, including cystic fibrosis, where dysregulated inflammatory responses contribute to disease progression. While air–liquid interface (ALI) models have advanced the study of airway epithelium, most of these modes lack immune components, limiting their ability to capture immune–epithelial interactions. Here, we expanded a previously established dual-cell ALI model incorporating human monocyte-derived macrophages to investigate how immune context, bacterial dose, and time influence early infection dynamics. Standard BCi-NS1.1 epithelial monocultures and macrophage co-cultures were infected with *P. aeruginosa* (PAO1) at low (100 colony-forming units (CFU) and high (1000 CFU) inoculum and analyzed over 10, 16, and 24 h post-infection (hpi). Macrophage presence did not significantly alter total bacterial burden but markedly influenced cytokine responses and bacterial spatial organization. Pro-inflammatory cytokines (interleukin (IL)-1α, IL-1β, Tumor Necrosis Factor (TNF)-α) were enhanced in dual-cell culture models, while IL-6 exhibited a threshold-dependent response detectable only at higher bacterial loads. Confocal imaging revealed that macrophages altered bacterial distribution, promoting a more dispersed pattern compared to the clustered organization observed in epithelial monocultures. These effects were most pronounced at lower bacterial inocula. Together, our findings demonstrate that macrophages reshape early infection dynamics by modulating inflammatory signaling and bacterial spatial organization without affecting overall bacterial burden. This study highlights the importance of incorporating immune cells into *in vitro* airway models.

## Introduction

The increasing prevalence of antibiotic-resistant bacterial infections represents a major challenge to global healthcare systems and underscores the need for physiologically relevant experimental models that more accurately recapitulate host–pathogen interactions (Wilke *et al*., 2011; Papazian, Würtzen and Hansen, 2016; McCarron, Donnelley and Parsons, 2018; Ahookhosh *et al*., 2020; McCarron, Parsons and Donnelley, 2021; Tran *et al*., 2022; Petpiroon *et al*., 2023). Among opportunistic pathogens, *Pseudomonas aeruginosa* is of particular clinical importance due to its intrinsic resistance to multiple antibiotics and its frequent association with hospital-acquired infections, ventilator-associated pneumonia and chronic respiratory infections (Hawdon *et al*., 2010; Fallows *et al*., 2016; Weber *et al*., 2016; Restrepo *et al*., 2018; Graf *et al*., 2023; Leoni Swart *et al*., 2024; Li *et al*., 2024).

In addition to its role in nosocomial infections, *P. aeruginosa* is a dominant pathogen in the lungs of people with cystic fibrosis (pwCF) where it establishes chronic infections that are difficult to eradicate. CF lung disease is characterized by impaired mucociliary clearance, leading to persistent bacterial infection and pronounced inflammation. A hallmark of CF airway disease is a dysregulated immune response, often associated with sustained production of pro-inflammatory cytokines such as interleukin-6 (IL-6), tumor necrosis factor-α (TNF-α), and IL-1 family members (IL-1α and IL-1β), which contribute to excessive neutrophil recruitment and progressive tissue damage. IL-1α primarily acts as an alarmin released following cellular stress or damage, whereas IL-1β and IL-6 are actively secreted in response to microbial stimulation (Boucher, 2007; Feliziani *et al*., 2014; Roussel *et al*., 2016; Bonfield and Chmiel, 2017; Berkebile *et al*., 2020; Ciszek-Lenda *et al*., 2023; Rossi *et al*., 2024).

Cytokine signaling plays a central role in coordinating host defense during airway infection by regulating immune cell recruitment and activation, and communication between epithelial and immune compartments. Early responses involve cytokines such as IL-1α and IL-1β, which initiate inflammatory signaling cascades, as well as IL-6, an important acute-phase cytokine that contributes to amplification of inflammatory responses and links innate and adaptive immunity. Chemokines such as monocyte chemoattractant protein-1 (MCP-1) further regulate immune cell recruitment (Fallows *et al*., 2016; He, Hara and Núñez, 2016; Totani *et al*., 2017; Roesch, Nichols and Chmiel, 2018). In addition, cytokines such as IL-10 and interferon-γ(IFN-γ) contribute to immune regulation through distinct mechanisms, with IL-10 functioning primarily as an anti-inflammatory mediator, while IFN-γ promotes macrophage activation and polarization of pro-inflammatory immune responses (Jäger *et al*., 2021; Paplinska-Goryca *et al*., 2021; Gopallawa *et al*., 2023). Dysregulation of these signaling pathways has been associated with chronic inflammation and impaired host defense in airway diseases (Shi and Pamer, 2011; Beale *et al*., 2014; Singh *et al*., 2015; Broz and Dixit, 2016; Kelley *et al*., 2019 Peña-Cearra *et al*., 2024).

Air–liquid interface (ALI) models have significantly advanced the study of respiratory infections by enabling differentiation of airway epithelial cells into pseudostratified structures that reproduce key features of the human airway, including mucus production, ciliary activity, and epithelial barrier formation (Laborda *et al*., 2024; Leoni Swart *et al*., 2024; Colque *et al*., 2026). However, conventional ALI models are typically limited to epithelial monocultures and lack immune components, restricting their ability to capture the dynamic interplay between epithelial and immune responses during infection (Melanson *et al*., 2026).

Recent studies have begun to incorporate immune cells into ALI systems to better reflect the airway microenvironment, demonstrating that immune–epithelial interactions can influence cytokine production and host responses to infection (Chikina *et al*., 2020; De Rudder *et al*., 2020; Plebani *et al*., 2022; Prescott *et al*., 2023; Burns *et al*., 2025; Colque *et al*., 2026). Despite these advances, important aspects of infection biology remain insufficiently explored and poorly understood. In particular, it remains unclear how immune cells modulate infection dynamics over time, how bacterial dose influences host responses, and how broader cytokine networks evolve during the early stages of infection.

In a previous study, we established a dual-cell ALI model incorporating human monocyte-derived macrophages on the basolateral side of differentiated airway epithelial cells, providing a platform to investigate host–pathogen interactions in a more physiologically relevant context (Melanson et al., 2026). Building on this model, the present study aimed to investigate how macrophage presence influences infection dynamics and cytokine responses across time and bacterial dose in non-CF epithelial cells. Although the model does not directly replicate CF airway disease, the inflammatory pathways investigated are relevant to both normal and diseased airway conditions. By expanding the temporal and immunological characterization of the airway model, this study provides new insight into how immune–epithelial interactions shape the early stages of airway infection.

## Materials and Methods

### Cell culture and establishment of the dual-cell ALI infection model

Polyester membrane transwell inserts with a pore size of 0,4 μm (Corning®) were used to establish the ALI culture system. The BCi-NS1.1 cell line, an immortalized human airway basal epithelial cell line capable of differentiating into a mucociliary epithelium under ALI conditions (Walters et al., 2013; Prescott et al., 2023), was used throughout the study. Cells were kindly provided by Ronald G. Crystal (Weill Cornell Medical College, New York, USA). BCi-NS1.1 cells were expanded and seeded onto collagen-coated transwell inserts and differentiated under ALI conditions for up to 28 days, as previously described (Laborda et al., 2024; Colque et al., 2026).

To promote epithelial attachment, inserts were coated apically with type I collagen (Gibco) and incubated for 30 min at room temperature. Inserts were subsequently inverted, and 20 µL of fibronectin (Merck; 5 µg/mL) was applied to the basolateral surface and incubated for 2 h at 37 °C. Fibronectin was diluted in phosphate-buffered saline (PBS, Gibco). Peripheral blood mononuclear cells (PBMCs) were isolated from buffy coats obtained from healthy donors (Department of Clinical Immunology, Rigshospitalet, Copenhagen, Denmark). Buffy coats from 3–4 donors were pooled per experiment to increase yield and reduce donor variability. PBMCs were isolated using LEUCOSEP separation tubes (Greiner) pre-filled with Lymphoprep (STEMCELL Technologies) following dilution in PBS supplemented with 2 % heat inactivated fetal bovine serum (FBS, Gibco). Cells were washed to remove platelets and cryopreserved at 1 × 10C cells/mL in FBS containing 10 % dimethyl sulfoxide (DMSO, Thermo Fisher Scientific).

For macrophage generation, PBMCs were thawed and cultured in M0 base medium consisting of RPMI 1640 supplemented with 10 % FBS and 50 ng/mL macrophage colony-stimulating factor (M-CSF, Miltenyi Biotec). Monocytes were isolated by negative selection (MojoSort™ Human Pan Monocyte Isolation Kit, BioLegend) and seeded at a density of 1.4 × 10^7^cells/mL. Cells were differentiated into macrophages over six days, with media replacement after 24 h to remove non-adherent cells. Prior to co-culture establishment, epithelial integrity was assessed by transepithelial electrical resistance (TEER) measurements. To enable macrophage adhesion, transwell inserts were inverted and coated with a secondary coating of fibronectin (5 µg/mL) on day 27 for 30 min.

Differentiated macrophages were seeded onto the basolateral surface of fully differentiated epithelial cultures. A volume of 20 µL macrophage suspension (8.5 × 10C cells) was applied and incubated for 1 h at 37 °C with 5 % COC to allow attachment. Control inserts were treated identically with macrophage-free medium. Following incubation, inserts were returned to their original orientation in 24-well plates containing 400 µL of pre-warmed medium consisting of a 1:1 mixture of M0 base medium and PneumaCult-ALI maintenance medium (STEMCELL Technologies), supplemented with heparin (4 µg/mL), hydrocortisone (480 ng/mL), PneumaCult-ALI 10× supplement, and PneumaCult-ALI maintenance supplement. Cultures were established and maintained as described previously (Melanson *et al.,* 2026)

### Preparation of bacterial inoculum and infection procedure

The laboratory strain *P. aeruginosa* PAO1 was cultured under standard conditions in Luria–Bertani (LB) medium. Bacterial inoculum was prepared based on colony forming unit (CFU) calculations. Infection of ALI cultures was performed apically using two defined inoculum sizes: 100 CFU and 1000 CFU. Infection experiments were conducted over multiple time points (10, 16, and 24 h) to assess infection kinetics. Both mono-cell (epithelial-only) and dual-cell (epithelial + macrophage) culture models were infected under identical conditions. The infection protocol was based on previously established methods (Laborda et al., 2024 & Melanson et al., 2026), with modifications to inoculum size and infection duration to accommodate the dual-cell configuration and enable analysis of infection dynamics over time.

### Quantification of bacterial burden and epithelial integrity

To assess bacterial distribution within the ALI model, three fractions were collected: apical-site, epithelial-associated, and basolateral-site. The apical fraction was obtained by washing the apical surface with PBS and represents planktonic and loosely associated bacteria. Epithelial-associated bacteria were recovered following treatment with Triton X-100, representing bacteria attached to or located within epithelial layers. The basolateral fraction was collected from the basolateral medium and represents bacteria that traversed the epithelial barrier. All samples were serially diluted and plated in triplicate on LB agar plates. CFUs were quantified and expressed as CFU/mL. Epithelial barrier integrity was assessed by measuring TEER using a voltohmmeter (e.g., EVOM2, World Precision Instruments) with STX/chopstick electrodes according to the manufacturer’s instructions. TEER values were corrected by subtracting blank insert resistance and normalized to membrane surface area.

### Cytokine responses

Cytokine secretion was quantified from basolateral fluids using a cytometric bead array (CBA) multiplex assay according to the manufacturer’s instructions (BD Biosciences, San Jose, CA, USA). The panel included IL-1β, IL-2, IL-6, IL-8, IL-10, IL-12p70, TNF-α, IFN-γ, and MCP-1 using the BD™ CBA Human Flex Set in combination with the BD™ Human Flex Set Master Buffer Kit (BD Biosciences).

Supernatants were collected at defined time points post-infection and stored at −80 °C until analysis. Samples were incubated with cytokine-specific capture beads and detection reagents according to the manufacturer’s specifications to form fluorescent bead–cytokine complexes. Data acquisition was performed using flow cytometry, and data were analyzed using BD CBA Analysis Software (version 1.1.15, BD Biosciences). Cytokine concentrations were determined using standard curves generated from recombinant cytokine standards provided in the assay kit.

### Immunostaining and confocal microscopy

Cells were fixed using 4% paraformaldehyde (PFA; Thermo Fisher Scientific). After fixation, samples were washed with PBS, and all subsequent staining and washing steps were performed directly within the transwell plate to preserve cellular architecture. For immunostaining, blocking and permeabilization were performed simultaneously using a buffer consisting of 1% (w/v) saponin (Sigma-Aldrich) and 3% (w/v) bovine serum albumin (BSA; Thermo Fisher Scientific) diluted in PBS and incubated for 1 h at room temperature. Primary antibodies were diluted in blocking buffer and incubated overnight at 4 °C. The following primary antibodies were used: anti-CD14 (Thermo Fisher Scientific) to identify macrophages and anti–acetylated α-tubulin (Santa Cruz Biotechnology) to visualize ciliated epithelial cells.

Following primary antibody incubation, membranes were washed three times with PBS in the plate. Secondary antibodies (goat anti-rabbit; Thermo Fisher Scientific) were diluted in blocking buffer and incubated for 1 h at room temperature protected from light. Nuclear staining was performed using DAPI (4′,6-diamidino-2-phenylindole; Thermo Fisher Scientific). In selected experiments, To-Pro-3 (Thermo Fisher Scientific) was used as an alternative nuclear stain to optimize spectral separation between fluorophores. After completion of staining, membranes were carefully excised from the transwell inserts using a sterile scalpel and mounted onto glass slides using VECTASHIELD® antifade mounting medium (Vector Laboratories). Coverslips were applied and sealed with nail polish.

Confocal imaging was performed using a Leica Stellaris 8 confocal microscope equipped with a 20× objective (NA 1.3). Z-stack images were acquired with a step size of 0.1 µm, and identical imaging settings (laser power, gain, and offset) were maintained across all experimental conditions. Z-stack images were processed using maximum intensity projection for visualization.

### Data analysis

Flow cytometry-based cytokine data were analyzed using BD CBA Analysis Software (version 1.1.15, BD Biosciences). Confocal images were processed using Fiji (ImageJ, version 2.14.0/1.54f), and Z-stack images were projected using maximum intensity projection in addition to this Z-stack sizes was 0.1µm.

Statistical analysis was performed using GraphPad Prism (version 10.5.0). Data are presented as mean ± standard deviation (SD) from three independent biological replicates, each with three technical replicates. Data distribution was assessed using the Brown–Forsythe test before statistical analysis. Statistical significance was determined using Tukey’s multiple comparisons test and two-way ANOVA with Tukey’s multiple comparisons test, with significance thresholds defined as *p ≤ 0.05, **p ≤ 0.01, ***p ≤ 0.001, and ****p < 0.0001.

## Results

### Epithelial barrier integrity is modestly affected by bacterial dose

TEER measurements revealed a dose-dependent reduction in epithelial barrier integrity at early time points, with significantly lower TEER values observed in models infected with 1000 CFU compared to 100 CFU at 10 h post-infection (hpi) (Figure 1A). However, no consistent differences in TEER were observed between mono-cell (BCi-NS1.1) and dual-cell (BCi-NS1.1 + macrophages) models under equivalent infection conditions, indicating that macrophage inclusion did not compromise epithelial barrier function. At later time points (16 and 24 hpi), TEER values remained relatively stable across conditions, with only minor variations observed (Figure 1A). CFU quantification at 10 and 16 hpi demonstrated that macrophage presence did not significantly alter total bacterial count across apical or epithelial-associated compartments when comparing mono- and dual-cell culture models at the same bacterial inoculum (Figure 1B–C). As expected, increasing the bacterial inoculum from 100 to 1000 CFU resulted in significantly higher bacterial counts, particularly in the apical fraction, in both mono- and dual-cell culture systems (Figure 1B–C). At 16 hpi, a modest increase in epithelial-associated bacteria was observed at higher inoculum, including a significant difference between mono- and dual-cell culture models under certain conditions, suggesting that macrophages may influence bacterial localization rather than overall CFU count (Figure 1C).

**Figure 1.**
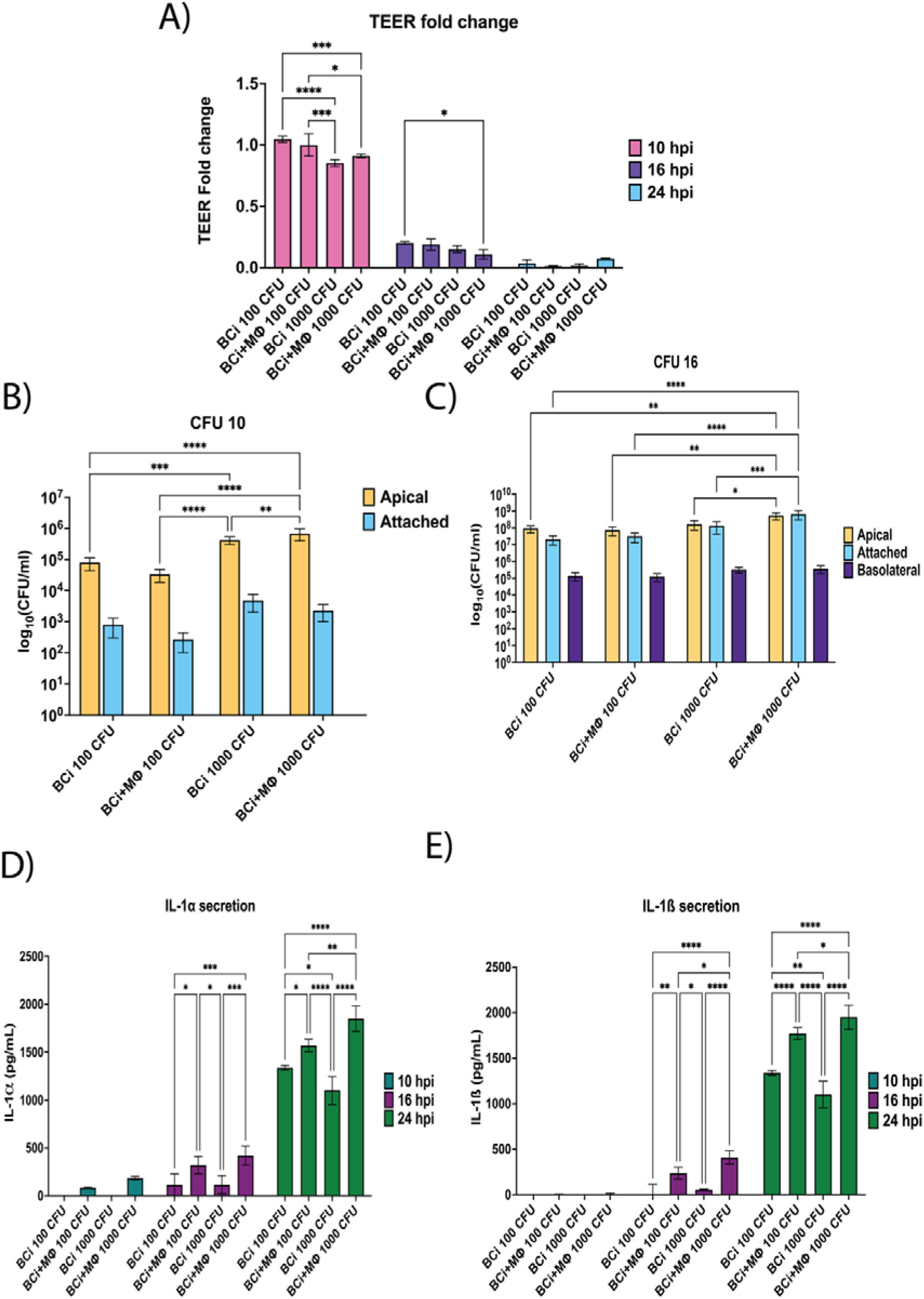
Infection dynamics and early inflammatory responses in mono-and dual-cell Air-Liquid Interface models. A) Transepithelial electrical resistance (TEER) measurements at 10, 16, and 24 h post-infection (hpi), showing epithelial barrier integrity in BCi-NS1.1 monocultures and BCi-NS1.1 + macrophage co-cultures infected with *Pseudomonas aeruginosa* (PAO1) at 100 or 1000 colony forming units (CFU). B) Quantification of apical bacterial burden (CFU/mL) at 10 hpi. C) Quantification of bacterial CFU/mL at 16 hpi, including apical and epithelial-associated fractions following Triton X-100 treatment. D) IL-1α secretion at 10, 16, and 24 hpi. E) IL-1β secretion at 10, 16, and 24 hpi. Apical fractions represent planktonic and loosely associated bacteria, while epithelial-associated fractions include bacteria attached to or internalized within epithelial cells. Cytokine concentrations were quantified using multiplex cytometric bead array (CBA). TEER values were corrected by subtracting blank insert resistance and normalized to membrane surface area. Data represent mean ± SD from three independent biological replicates, each with three technical replicates. Statistical significance was determined using two-way ANOVA with Tukey’s multiple comparisons test (*p ≤ 0.05, **p ≤ 0.01, ***p ≤ 0.001, ****p < 0.0001).

Analysis of IL-1α concentration showed minimal or absent IL-1α release in epithelial monocultures at 10 hpi, whereas dual-cell culture models exhibited detectable IL-1α levels even at low bacterial inoculum (100 CFU) (Figure 1D). At 16 and 24 hpi, IL-1α levels increased in both models, with significantly higher release observed in macrophage-containing cultures, particularly at higher bacterial doses (Figure 1D). A similar trend was observed for IL-1β. No detectable IL-1β secretion was observed at 10 hpi, while increased IL-1β levels were detected at 16 and 24 hpi, predominantly in dual-cell culture models (Figure 1E). These increases were significantly higher compared to epithelial monocultures and were further enhanced at 1000 CFU.

### Macrophages reshape the cytokine landscape in a dose- and time-dependent manner

To further characterize immune signaling in the model, cytokine secretion was assessed at 10, 16, and 24 hpi in mono- and dual-cell ALI cultures (Figure 2). IL-2 displayed a transient profile, with detectable levels at 10 hpi followed by a reduction at 16 hpi and no detectable IL-2 at 24 hpi (Figure 2A). At 1000 CFU and 10 hpi, IL-2 levels were higher in dual-cell cultures compared to epithelial monocultures, whereas at later time points IL-2 levels declined across conditions. A distinct threshold-dependent response was observed for IL-6, which was only detectable at the higher inoculum (1000 CFU) across all time points (Figure 2B). IL-6 levels were elevated in both epithelial monocultures and dual-cell cultures at 1000 CFU, with higher levels observed in dual-cell cultures at early time points. At 24 hpi, epithelial monocultures also contributed substantially to the total IL-6 response, indicating that IL-6 production under high bacterial burden conditions is not restricted to macrophage-containing cultures.

**Figure 2.**
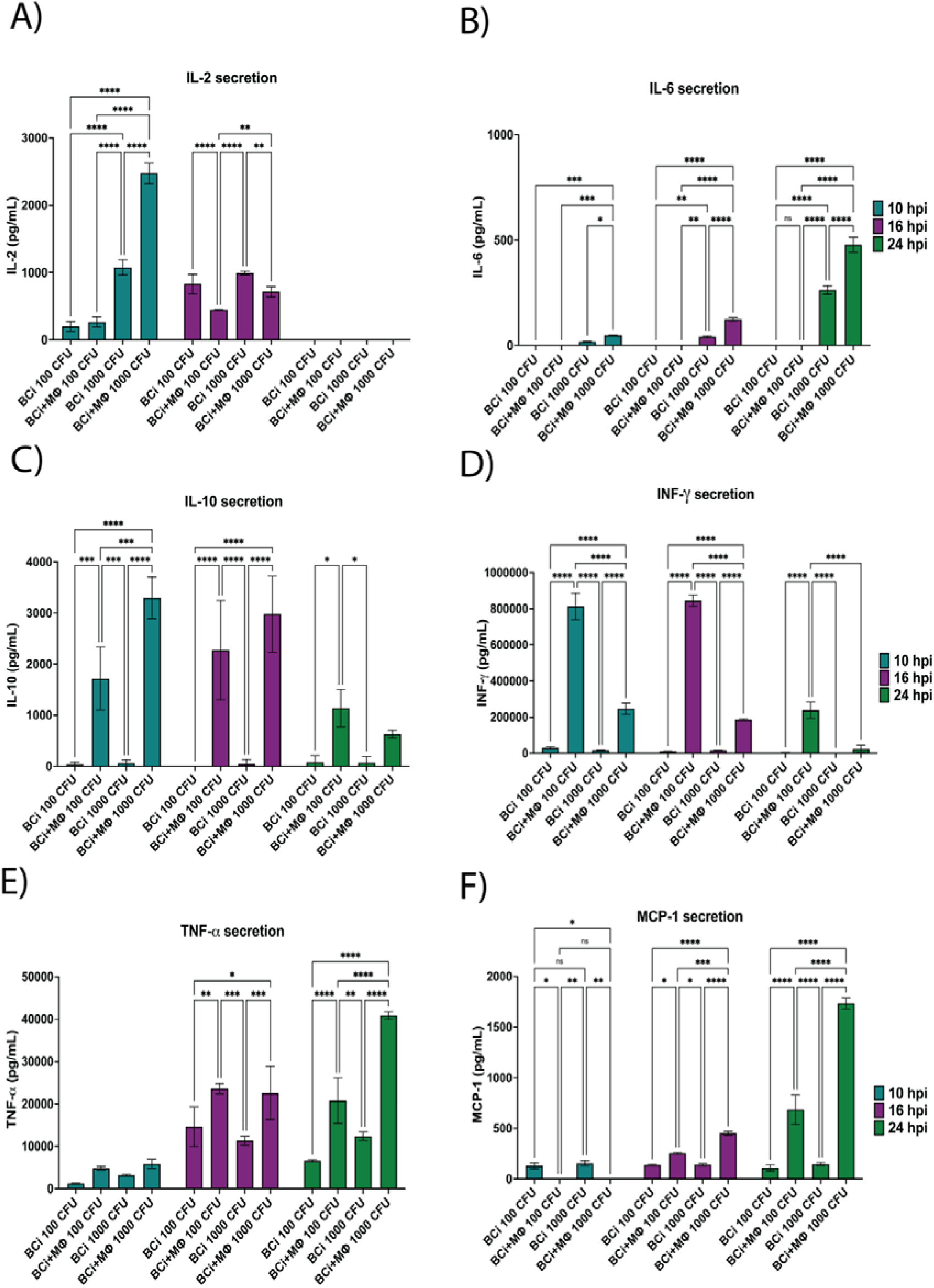
Cytokine profiling of mono- and dual-airway cell Air-Liquid Interface models during infection. Cytokine secretion was measured at 10, 16, and 24 h post-infection (hpi) with *Pseudomonas aeruginosa* (PAO1) at 100 or 1000 colony forming units (CFU) in BCi-NS1.1 monocultures and BCi-NS1.1 + macrophage co-cultures. A) Interleukin (IL)-2 secretion, B) IL-6 secretion, C) IL-10 secretion, D) Interferon-γ (IFN-γ) secretion, E) Tumor necrosis factor-α (TNF-α) secretion, F) Monocyte chemoattractant protein-1 (MCP-1) secretion. Cytokine concentrations were quantified using multiplex cytometric bead array (CBA). Data represent mean ± SD from three independent biological replicates, each with three technical replicates. Statistical significance was determined using Two-way ANOVA with Tukey’s multiple comparisons test and is indicated as *p ≤ 0.05, **p ≤ 0.01, ***p ≤ 0.001, ****p ≤ 0.0001.

IL-10 and IFN-γ displayed distinct response patterns and were therefore considered separately. IL-10 was primarily detected in macrophage-containing cultures and was highest at earlier time points, with reduced levels observed over time under high-inoculum conditions (Figure 2C). In contrast, IFN-γ was also mainly detected in macrophage-containing cultures, but high bacterial inoculum was associated with lower IFN-γ levels across time points (Figure 2D). These findings indicate that IL-10 and IFN-γ are differentially regulated in response to bacterial dose and time. At later time points, TNF-α and MCP-1 levels increased, particularly in dual-cell culture models (Figure 2E–F). TNF-α secretion was enhanced in macrophage-containing cultures at 16 and 24 hpi, while MCP-1 showed limited levels at early time points but increased over time, especially in the presence of macrophages. Together, these data show that macrophage-containing cultures reshape cytokine responses in a dose- and time-dependent manner.

### Macrophages alter the spatial organization of *P. aeruginosa* in a time and dose dependent manner

To further investigate how macrophages influence infection dynamics beyond total bacterial count, confocal imaging was performed to assess the spatial organization of *P. aeruginosa* in mono- and dual-cell ALI models (Figure 3). At 10 hpi with 100 CFU, epithelial monocultures displayed localized bacterial clustering at the apical surface, consistent with early aggregate formation (Figure 3A). In contrast, dual-cell culture models containing macrophages exhibited a more dispersed bacterial distribution, with bacteria appearing more evenly distributed across the epithelial surface. These observations are consistent with previously reported findings under similar conditions (Melanson et al., 2026). Increasing the bacterial inoculum to 1000 CFU resulted in increased bacterial density and more pronounced clustering in both mono- and dual-cell culture models at 10 hpi (Figure 3B). To support the qualitative observations from confocal imaging, bacterial cluster area was quantified at 10 hpi following infection with 100 CFU (Figure 4A). GFP-positive bacterial aggregates in epithelial monocultures displayed significantly larger cluster areas compared with dual-cell culture models containing macrophages. In contrast, macrophage-containing cultures exhibited significantly smaller and more dispersed bacterial clusters, supporting the observation that macrophages alter bacterial spatial organization during early infection.

**Figure 3.**
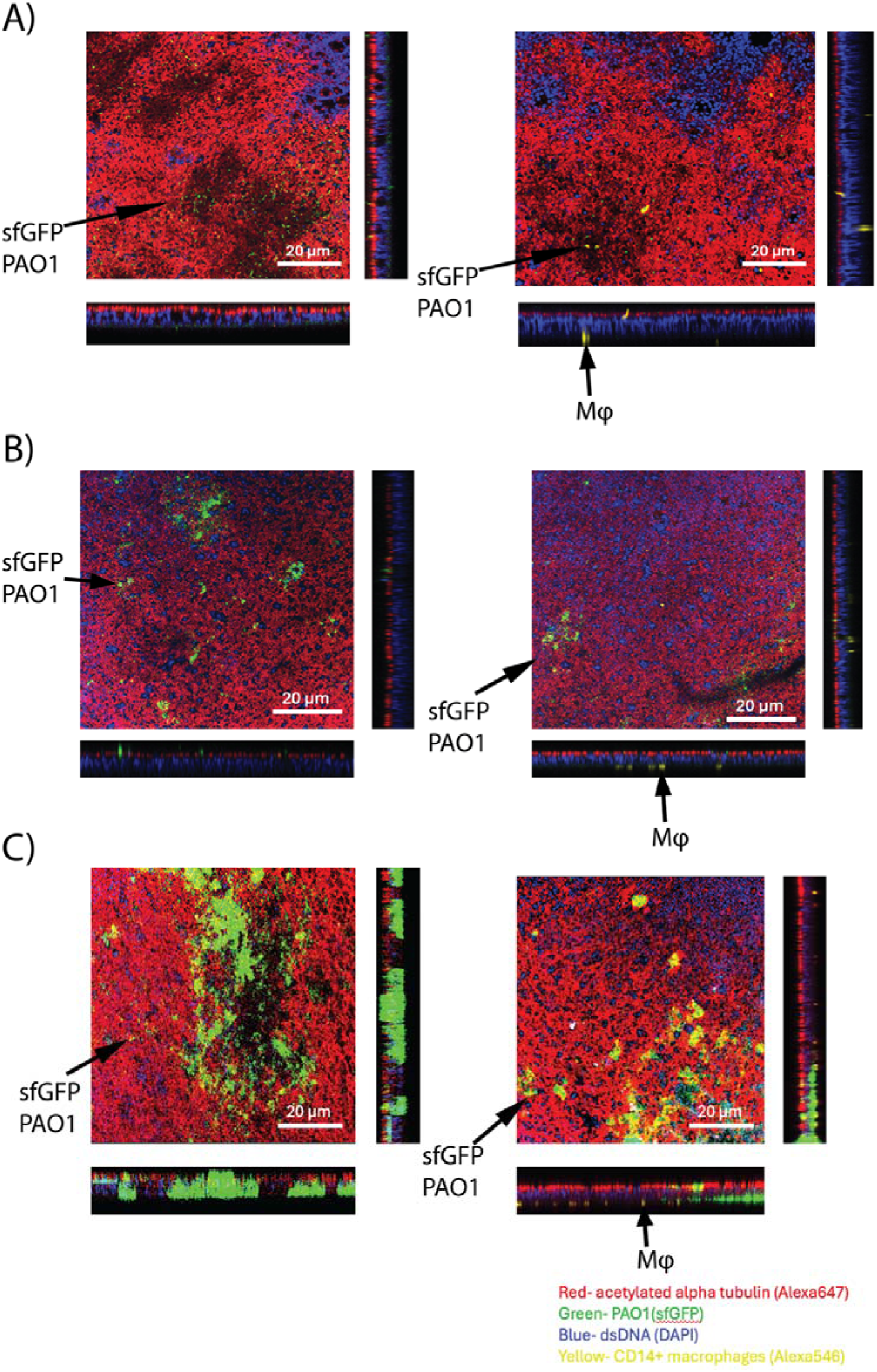
Macrophages modulate the spatial organization of *Pseudomonas aeruginosa* in the Air-Liquid Interface model. Confocal images of *Pseudomonas aeruginosa* (PAO1, sfGFP-expressing) infection in BCi-NS1.1 monocultures and BCi-NS1.1 + macrophage co-cultures under varying infection conditions. A) Infection with 100 colony forming units (CFU) at 10 h post-infection (hpi), showing localized bacterial clustering in epithelial monocultures and a more dispersed bacterial distribution in dual-cell culture models. B) Infection with 1000 CFU at 10 hpi, demonstrating increased bacterial density and clustering in both models, with a more heterogeneous distribution maintained in dual-cell cultures. C) Infection with 1000 CFU at 16 hpi, showing extensive bacterial accumulation and loss of defined spatial organization, with regions of heterogeneous distribution and macrophage-associated interactions in dual-cell culture models. Epithelial cells are visualized by acetylated α-tubulin staining (red, Alexa Fluor 647), bacteria are shown as GFP-expressing PAO1 (green), nuclei are stained with DAPI (blue), and macrophages are identified by CD14^+^ staining (yellow, Alexa Fluor 546). For each condition, top (en face) views are shown as maximum intensity projections, alongside corresponding orthogonal (x–z and y–z) views, illustrating the vertical distribution of bacteria relative to the epithelial layer. Arrows indicate representative macrophages and bacterial clusters within the images. Images were acquired using a confocal microscope at 20× magnification. Identical acquisition settings were applied across all conditions. Scale bars: 20 µm.

**Figure 4.**
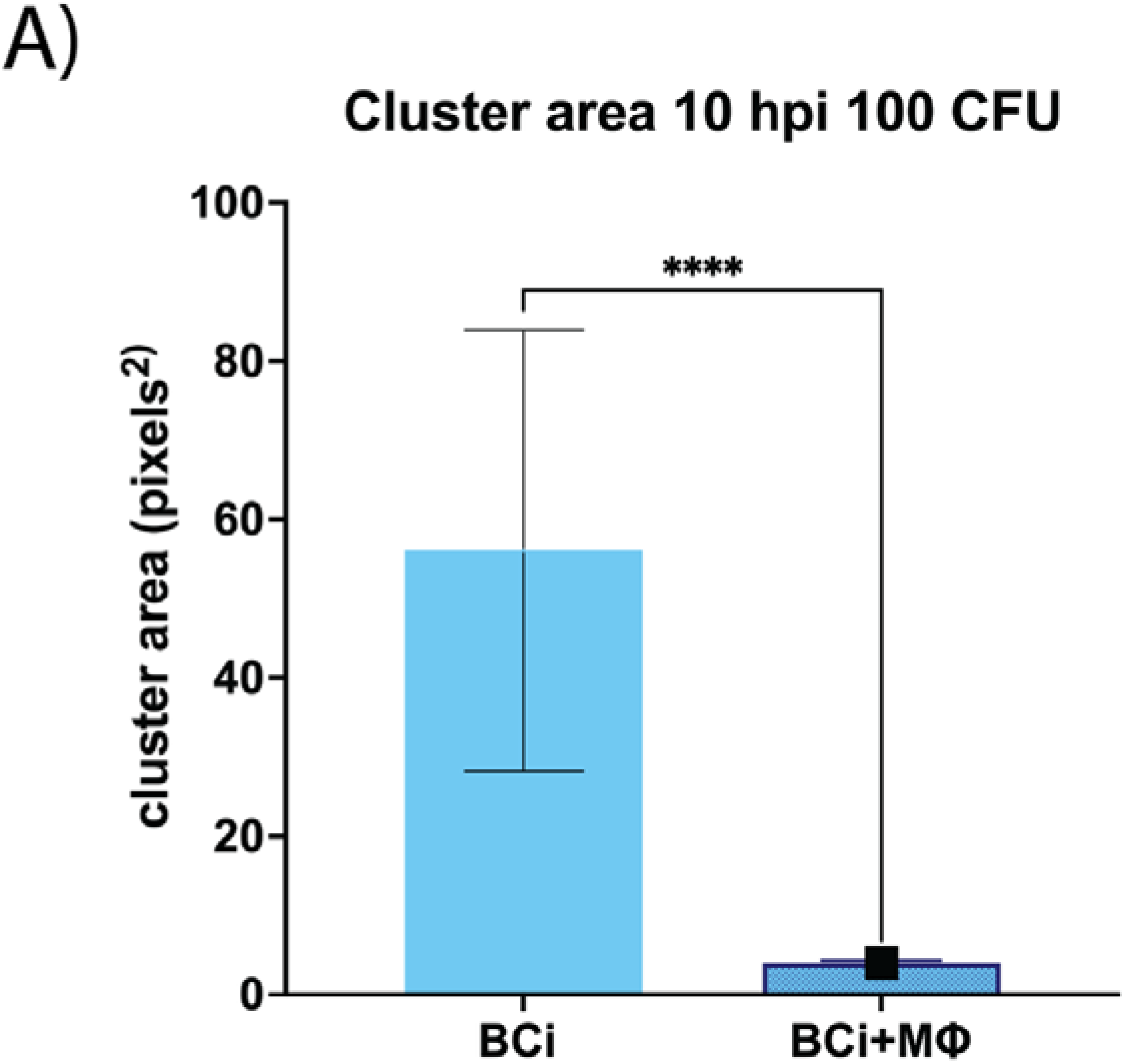
Quantification of bacterial cluster area in mono and dual-cell Air-Liquid Interface models at early infection stages. A) Quantification of GFP-positive bacterial cluster area at 10 h post-infection (hpi) following infection with Pseudomonas aeruginosa (PAO1) at 100 CFU in BCi-NS1.1 monocultures and BCi-NS1.1 + macrophage co-cultures. Cluster area was measured from confocal images using Fiji/ImageJ and expressed as pixels². Macrophage-containing co-cultures exhibited significantly smaller bacterial cluster areas compared with epithelial monocultures, consistent with a more dispersed bacterial distribution. Data represent mean ± SD from three independent biological replicates, each with three technical replicates. Statistical significance was determined using an unpaired t-test with Welch’ s correction (****p < 0.0001).

At 16 hpi with 1000 CFU, bacterial overgrowth and aggregation were observed in both models, with extensive surface coverage and loss of clearly defined spatial organization (Figure 3C). Orthogonal projections confirmed that bacteria were predominantly localized at the apical surface at early time points, while increased signal intensity and broader distribution were observed across the epithelial layer at higher bacterial loads and later time points (Figure 3A–C). Importantly, these differences in spatial organization occurred despite no significant differences in total bacterial count between mono- and dual-cell culture models (Figure 1B–C), indicating that at early time points of infection, macrophages influence bacterial distribution and infection architecture rather than overall bacterial numbers.

## Discussion

This study expands upon our previously established dual-cell culture ALI model to investigate how immune cells, bacterial dose, and time influence early infection dynamics in the airway epithelium. By incorporating macrophages and examining infection across multiple time points and inoculum sizes, we demonstrate that host responses and bacterial behavior are modulated by both bacterial inoculum and immune–epithelial interactions.

Infection progression was characterized by changes in epithelial integrity and cytokine production over time, while total bacterial burden remained relatively stable, suggesting that early infection dynamics in this model are primarily driven by host responses rather than bacterial expansion. Macrophage-containing cultures exhibited enhanced cytokine responses compared with epithelial monocultures, including increased levels of IL-1 cytokines, TNF-α, and MCP-1, supporting a role for macrophages in shaping the inflammatory microenvironment during early infection. Although this model is not intended to directly replicate chronic airway infection, it captures early epithelial–immune interactions that may influence subsequent inflammatory progression, immune cell recruitment, and bacterial persistence. These findings provide insight into early host–pathogen interactions that may contribute to later disease development and therapeutic responses.

The observed increase in cytokine secretion over time is consistent with previous studies demonstrating accumulation of bacterial and host-derived inflammatory mediators during *P. aeruginosa* infection (Singh *et al*., 2015; Jäger *et al*., 2021; Blevins *et al*., 2022; Ciszek-Lenda *et al*., 2023). In the present study, cytokine responses displayed clear dose-dependent patterns. IL-6 secretion was only detected at the higher bacterial inoculum, suggesting that activation of this cytokine requires a threshold level of bacterial stimulation. Similar dose-dependent IL-6 responses have previously been associated with activation of acute inflammatory signaling pathways during bacterial infection (Dillingh *et al*., 2014). In contrast, IL-1α, IL-1β, and TNF-α were induced at both low and high bacterial inocula and were consistently enhanced in macrophage-containing cultures. These findings suggest that macrophage presence contributes to amplification of inflammatory signaling within the epithelial co-culture environment rather than simply reflecting bacterial burden alone (Esvaran and Conway, 2018). Together, these observations indicate that macrophages selectively modulate early inflammatory responses and that bacterial inoculum influences both the magnitude and character of airway immune activation.

Our findings are consistent with previous studies demonstrating that airway epithelial cells are important sources of chemokines and pro-inflammatory cytokines during *P. aeruginosa* infection. Bacterial components such as flagellin and lipopolysaccharide activate epithelial innate immune signaling pathways, leading to production of cytokines including IL-6, IL-8, and IL-1 family members (Knowles and Boucher, 2002; Kawai and Akira, 2010; Whitsett and Alenghat, 2015; Yin *et al*., 2015; Bastaert *et al*., 2018; Berkebile *et al*., 2020; Acosta and Alonzo, 2023; Graf *et al*., 2023; Norris and Kubes, 2025). In addition to direct epithelial signaling, macrophages are known to amplify local inflammatory responses through cytokine-mediated communication with epithelial cells and recruitment of additional immune populations. The increased cytokine secretion observed in the dual-cell culture model therefore supports the importance of epithelial–immune interactions during early airway infection. Although IL-12p70 was not detected in the present study, IFN-γ secretion was still observed, suggesting activation through IL-12-independent pathways or contribution from alternative signaling mechanisms within the co-culture system (Wilke *et al*., 2011; Bonfield and Chmiel, 2017; Roesch, Nichols and Chmiel, 2018; Qionghua Chen, Yuelin Shen and Jingyang Zheng, 2021; Møller *et al*., 2024; Rossi *et al*., 2024; Colque *et al*., 2026).

Previous *in vitro* studies using airway epithelial monocultures have shown that *P. aeruginosa* infection induces early production of cytokines such as IL-6 and IL-8, followed by later induction of IL-1β and TNF-α (Denning *et al*., 1998; Wu *et al*., 2005; Zhang, Wu and Yu, 2005; Shanks *et al*., 2010). In contrast, studies using CF-derived epithelial models and more advanced co-culture systems have demonstrated enhanced and sustained inflammatory responses, reflecting the dysregulated immune environment associated with chronic CF airway disease (Totani *et al*., 2017; Plebani *et al*., 2022; Cadenas-Jiménez *et al*., 2025). Notably, Plebani et al. demonstrated persistent inflammatory activation in a CF airway-on-chip model following *P. aeruginosa* infection, including elevated cytokine production and altered epithelial responses. Compared with these CF-associated models, the cytokine profiles observed in the present study were more regulated and strongly dependent on bacterial inoculum and time. This likely reflects the use of non-CF epithelial cells and supports the interpretation that the present model captures early-stage airway immune responses prior to the development of chronic inflammatory dysregulation. In this context, macrophage-mediated modulation of cytokine signaling may represent early regulatory mechanisms that become altered or amplified during chronic CF infection.

Despite similar bacterial burdens between conditions, confocal imaging demonstrated clear differences in bacterial spatial organization depending on macrophage presence. In epithelial monocultures, *P. aeruginosa* formed larger and more localized bacterial aggregates, whereas bacteria in dual-cell culture models appeared smaller and more dispersed. Quantification of bacterial cluster area supported these qualitative observations and demonstrated significantly reduced aggregate size in macrophage-containing cultures. These findings suggest that macrophages influence the local airway microenvironment in ways that alter bacterial organization without substantially affecting total bacterial numbers. Similar concepts have previously been proposed in studies showing that immune cells can influence bacterial behavior through modulation of cytokines, antimicrobial peptides, and mucus-associated factors rather than through direct bacterial clearance alone (Raoust *et al*., 2009; Farias and Rousseau, 2016; Bustamante-Marin and Ostrowski, 2017; Evren *et al*., 2022; Basil *et al*., 2024; Fu *et al*., 2025; Sahu and Ruhal, 2025).

The effect of macrophages on bacterial spatial organization was strongly influenced by bacterial inoculum. At lower infection doses, macrophage-dependent modulation of bacterial distribution was clearly observed, whereas higher inocula partially restored the aggregated growth pattern seen in epithelial monocultures. These findings suggest that increasing infection pressure may override early immune-mediated effects on bacterial organization. In chronic airway infections such as CF, *P. aeruginosa* progressively transitions from dispersed colonization toward more structured aggregates and biofilm-like growth that are difficult to eradicate (Jørgensen *et al*., 2024). The more dispersed bacterial phenotype that we observed in macrophage-containing cultures at lower inoculum may therefore reflect early host-mediated restriction of aggregate formation, whereas restoration of clustered growth at higher bacterial loads may resemble early progression toward persistent infection states. Together, these findings support the idea that early epithelial–immune interactions may contribute to shaping bacterial spatial organization during the initial stages of airway colonization.

The static transwell system is a limitation since it does not fully reproduce physiological airway conditions such as airflow, mucus clearance, and continuous nutrient exchange. Despite this, the model system provides a controlled and physiologically relevant platform for investigating early host–pathogen interactions and demonstrates that macrophages can substantially influence cytokine responses and bacterial spatial organization during early airway infection. By incorporating immune components and examining infection across time and dose, this study highlights the importance of immune–epithelial interactions in shaping infection outcomes and provides a framework for future mechanistic studies of airway infection dynamics.

## Ethical approval and consent

This study was approved by the local ethics committee of the Capital Region of Denmark (Region Hovedstaden), registration number H-20024750.

## Conflict of Interest

The authors declare that the research was conducted in the absence of any commercial or financial relationships that could be construed as a potential conflict of interest.

## Authors contribution

AFM, JJP, SM and HKJ contributed to the conception and design of the study. AFM performed the experimental work and statistical analysis. AFM, SM and HKJ wrote the manuscript. All authors contributed to manuscript revision, read, and approved the submitted version.

## Funding

The work at the Novo Nordisk Foundation Center for Biosustainability is supported by a grant from the Novo Nordisk Foundation (Ref. nr.: NNF10CC1016517). HKJ was supported by a Challenge Grant NNF19OC0056411 from The Novo Nordisk Foundation, and a grant from The John and Birthe Meyer Foundation. HKJ and SM were supported by a grant from CAG - Greater Copenhagen Health - Science - Partners 2020 (GCHSP).

## Acknowledgements

The authors thank the staff at the blood bank at Rigshospitalet (Copenhagen, Denmark) for preparing of the buffy coats for this study. Furthermore, the authors thank Ronald G. Crystal (Weil Cornell Medical College, New York, USA) for providing the BCi-NS1.1 cell line.

## Notes

### Competing Interest Statement

The authors have declared no competing interest.

